# Rules of departure: Antiphony and personalized vocal spaces in wild male elephant group coordination

**DOI:** 10.1101/2024.02.07.579388

**Authors:** Emmanouela Rantsiou

**Affiliations:** School of Engineering and Applied Sciences, Harvard University, Cambridge, MA, USA

## Abstract

In the intricate realm of animal communication, vocal signals play a vital role in maintaining bonds and coordinating group activities. Through detailed analysis of low-frequency rumble vocalizations of wild male African Savannah elephants, we demonstrate how these calls are employed in a coordinated manner. Our findings reveal a distinct pattern of call-and-response, usually initiated by a dominant male, with subsequent turn-taking responses from other group members. This antiphonal calling is instrumental in facilitating collective decision-making and group departures, highlighting a sophisticated level of communication previously unrecognized in male elephant groups. Furthermore, the study delves into the frequency content of these rumbles, finding significant individual variations in vocal characteristics. Analysis of the first two formant frequencies of these calls uncovers a distinct vocal space for each elephant, variable over time and across different social contexts. These individualized vocal spaces suggest a mechanism for personal identification within and across elephant groups, and may also relate to the information content of their vocal exchanges. The vocal spaces and structured antiphony presented in this study underline the complexity of elephant vocal communication and invite further targeted research into the vocal communication system of elephants, as well as inquiry into the cognitive and social aspects of wildlife communication in general.

## 1 Introduction

In social animals, one of the functions of vocal signaling is to maintain group cohesion by coordinating movements and departure [1–6], thus facilitating the benefits of group-living and preventing individuals from becoming unintentionally separated from their collective safety net. So-called ‘pre-departure vocalizations’ are often part of this coordinating and decision-making procedure. Such vocalizations have been reported in various species, including squirrel monkeys [7], hamadryas baboons [8], white-faced capuchin monkeys [9, 10], red-fronted lemurs [11], gorillas [12], swans [13], wild dogs [14], and elephants [15]. It is thought that in many cases, the use of repeated vocalizations prior to departure helps to reach a decision threshold - something like a voting mechanism - which triggers action [5, 6]. Departure vocalizations also serve to indirectly alert others at a distance to the fact that some particular area, e.g. a waterhole, may soon become available as a resource [16].

In the field of animal communication, the African Savannah elephant (Loxodonta Africana) with its distinct social structures and richness of vocal repertoire, stands as a captivating subject of inquiry. Elephants form various complex social groups [17], including i) family units composed of related females and their young offspring [18, 19], ii) bond groups and clans in which multiple families or groups of families coalesce into - usually temporary - social entities [18–21], and iii) small all-male groups [19, 22].

Elephants exhibit a broad array of vocalizations, encompassing the snort, trumpet, croak, chuff, rev, etc., and various types of rumble calls [15, 18, 23–25]. They employ a particular type of rumble, the “let’s go” rumble (LGR) [15], within families and bond groups in order to maintain cohesion and spatial coordination. These rumbles comprise a series of coordinated calls usually initiated by the matriarch or another dominant female to signal group departures [16,26]. This vocal coordination has the form of antiphonal calling.

The research on antiphonal calls in animals has seen an increase in attention and publications over the last decade [27]. True antiphonal vocalizations, also called cooperative vocal turn-taking, have been reported in the scientific literature - other than in humans - in a few species including apes [28, 29], monkeys [30–34], and a few other primates, cetaceans [35–37], [38], meerkats [39], and elephants [16, 40, 41] (though only in families and not in male groups; but see [42] for anecdotal reporting). A comprehensive overview of antiphonal calls across species and taxa can be found in [43].

Turn-taking in vocal communication is considered a key characteristic of linguistics [44] and has attracted special interest in research of language evolution. It has been shown (in humans [45] and non-human primates [31]) that turn-taking appears earlier than language skills in ontogeny and is learned through parental guidance. Understanding the origin, functionality, and usage of the turn-taking system of vocalization across species through comparative studies, can contribute to the understanding of language evolution [31,43]. Additionally, it provides an insight into cognitive processes related to speech comprehension and processing [46].

Studies on captive female African Savannah elephants have demonstrated the use of antiphonal vocalizations in affiliated individuals in a variety of social contexts [40]. In that study it was reported that these calls were clustered in bouts of rumbles separated by periods of relevant vocal inactivity. Note that, such clustering of elephant rumble callings has been previously reported on wild and captive Asian and African elephants [15, 24, 47]. In [41], the authors were able to further explore the functionality of these antiphonal rumble calls, and concluded that these temporarily associated rumbles were true communicative events, rather than common responses to shared external stimuli. They opined that the functionality of these antiphonal exchanges includes coordination of the herd’s movements and reinforcement of social bonds between herd members during reunions. Subsequent studies of female elephant families, this time in the wild, present a similar picture: cooperative antiphonal vocal exchanges are used among multiple family members in a communicative fashion, in order to facilitate group cohesion and coordination during departures from a watering ground [16].

The study of elephant vocalizations, in a variety of settings and contexts, has been focused predominantly on female individuals and families, while less attention has been given to male elephant groups, as males appeared significantly less active vocally. Targeted studies on male elephants however, are illuminating a yet undiscovered aspect of their vocal repertoire [48–50]. Our current study also focuses on male African elephant vocalizations - rumbles in particular.

The aim of this paper is twofold: 1) to document for the first time the use of antiphonal calls - in the form of rumbles - within groups of bonded male L. africana elephants, 2) to explore the vocal space of these rumble calls - as characterized by their first two formant frequencies - searching for vocal signatures at the level of individual elephants or group vocal events.

In the ‘Results’ section we showcase for the first time evidence of coordinated antiphonal rumble calls in groups of male African Savannah elephants (§2.1). We present the frequency characteristics of these calls (§2.2) and investigate the vocal spaces of individual elephants in and across group vocalization events and through the passage of time (§2.3). In the ‘Discussion’ section we make comparisons between the coordinated vocalization events presented and those reported previously in families of African Savannah elephants, and conclude that male elephant groups similarly use such rumble calls for social spatial coordination. We draw parallels between the antiphonal calls of the elephants to the turn-taking system of vocal exchanges in humans and other species. We also discuss the findings of our statistical analysis of vocal spaces and how individuality, social relations and interactions, or information content of rumble calls might contribute to these findings. The ‘Methods’ section contains further information on the dataset used for this study, the data reduction and analysis. Further information on some of the statistics tools we have used is also included. Finally, ‘Supplementary’ material is available with some extra figures and a link to online repository with the reduced data used in this study.

## 2 Results

The recordings of low frequency rumbles of male L. Africana took place in the wild over the period of 12 years (2005-2017), in the context of daily group visits at the Mushara waterhole in Etosha National Park, Namibia. The dataset includes six waterhole visit events of different duration and varying composition and size of the male elephant group. Some individuals appear in more than one of these visits. Table 1 and §4 (‘Methods’) provide further details on the data and each of these recorded events.

**Table 1:** Summary of rumble antiphonal exchange events used in this paper.

| **Event ID** | **05** | **07AS** | **07A** | **07B** | **11A** | **11BS** | **11B** | **17** |
| --- | --- | --- | --- | --- | --- | --- | --- | --- |
| *N* callers | 4 | 1 | 5 | 3 | 3 | 1 | 5 | 4 |
| *N* group members | 4(?) | 5 | 5 | 4 | 4(?) | 8 | 8 | 4 |
| *N* rumbles | 20 | 10 | 24 | 8 | 11 | 6 | 25 | 20 |
| *N* bouts | 8 | – | 9 | 3 | 2 | – | 8 | 8 |
| Average rumble duration [s] | 4.3 | 4 | 4.5 | 4.7 | 4.8 | 6 | 6.1 | 4.9 |
| Average bout duration [s] | 10.4 | – | 10.4 | 10.4 | 17.85 | – | 13.7 | 14.3 |
| Average time between rumbles [s] | 57.8 | 99 | 25.8 | 38.2 | 24.9 | 138.2 | 31.5 | 31.8 |
| Average time between bouts [s] | 157.9 | – | 353.5 | 138 | 51 | – | 81.3 | 98.3 |
| Median time between rumbles [s] | 0.9 | 44 | 2 | -1 | 1.35 | 79 | 0.8 | 2.1 |
| Median time between bouts [s] | 126 | – | 71 | 138 | 57 | 41.5 | 109 | 75.5 |
| Duration of visit [min] | 49 | – | 71 | – | 32 | – | 43.5 | – |
| Time between 1st rumble and departure [min] | 24 | – | 29 | – | 5 | – | 13.5 | – |
| Average F1 [Hz] | 23±3.7 | 25.2±2.2 | 28.3±2.7 | 27.2±4.3 | 21.8±2.4 | 18.5±1.4 | 21.1±4.2 | 18.8±2.7 |
| Average F2 [Hz] | 78.9±6.9 | 79.2±6.6 | 74.1±5.6 | 71.6±3.6 | 80.4±4.6 | 82.1±4.9 | 81.4±4.7 | 87±6.4 |

All calls considered in this study are low frequency rumbles. We observe three types of rumbles: i) single, isolated, and presumably unanswered rumbles uttered predominantly by the highest ranking elephant, ii) rumbles in bouts, i.e. sequences of rumbles with very short inter-call intervals (*<* 2*s*), iii) rumbles in bouts, but with partial overlap, i.e. negative, but short inter-call duration.

### 2.1 Antiphonal calls

We start by examining the vocal dynamics of these waterhole visit events in terms of number and duration of vocalizations, identity of the callers, timing of vocalizations, and overall time development of the sequence of vocalizations through the event. Our aim is to determine if the male African Savannah elephants use these types of rumbles as a way to coordinate their actions; in this case their departure from the watering ground.

Fig 1 shows in detail how event 07A unfolded, in time and space. A group of five elephants arrived at the waterhole, in the late evening. The recording was initiated upon arrival of the elephant group at the waterhole. Single rumbles as well as bouts (i.e. sequences) of rumbles are shown and marked accordingly. All ‘silent’ intervals between vocalizations have been removed from the main part of the plot for ease of visualization, but the bar at the lower part of the plot contains to scale the time intervals between vocalizations. Also in the lower bar, we encode in color the distance from the water trough where Greg - the group’s dominant bull - was, during the event. The whole event lasted ∼6000s. The vocalizations begin with a single rumble after 1890s of silence, followed by a bout of two overlapping rumbles, 683s later. For the following 2625s (Part A) a series of single, isolated rumbles take place, averaging 1rumble/180s (median: 1rumble/47s). This is a lower limit on the rate of single rumbles for this part of the event, as we only accounted for rumbles that were clearly identifiable in spectrograms. More single rumbles might have been present, in addition to the 14 rumbles presented in the figure. We note that 10 of those were identified as being produced by Greg; we do not know who the caller(s) of the others were, but it is possible that they were also produced by Greg. During this period of single rumbles, Greg is staying at the head of the trough, but starts moving away from it - at 50m distance - as he keeps uttering a few more single unanswered rumbles. This initial series of single rumbles is then followed by a more vocally active period of approximately 750s (Part B). We observe a number of rumble bouts interspersed with few single, isolated rumbles. Five individuals, including Greg, have been identified partaking in these rumble bouts. During this period, the rate or rumbles in bouts is 1rumble/28s (with a median of 1rumble/2s) and the average rate of all rumbles, single and in bouts, is 1rumble/31s (median of 1rumble/3.2s). By the time the rest of the group had begun engaging in bouts of rumbles - around the third bout, Greg had moved further away to 120m from the trough. The end of the rumble bouts finds Greg together with most of the group exiting the clearing, at 300m from the trough. A couple of single rumbles were uttered as the elephants were seen moving as a group away from the watering grounds.

**Figure 1:**
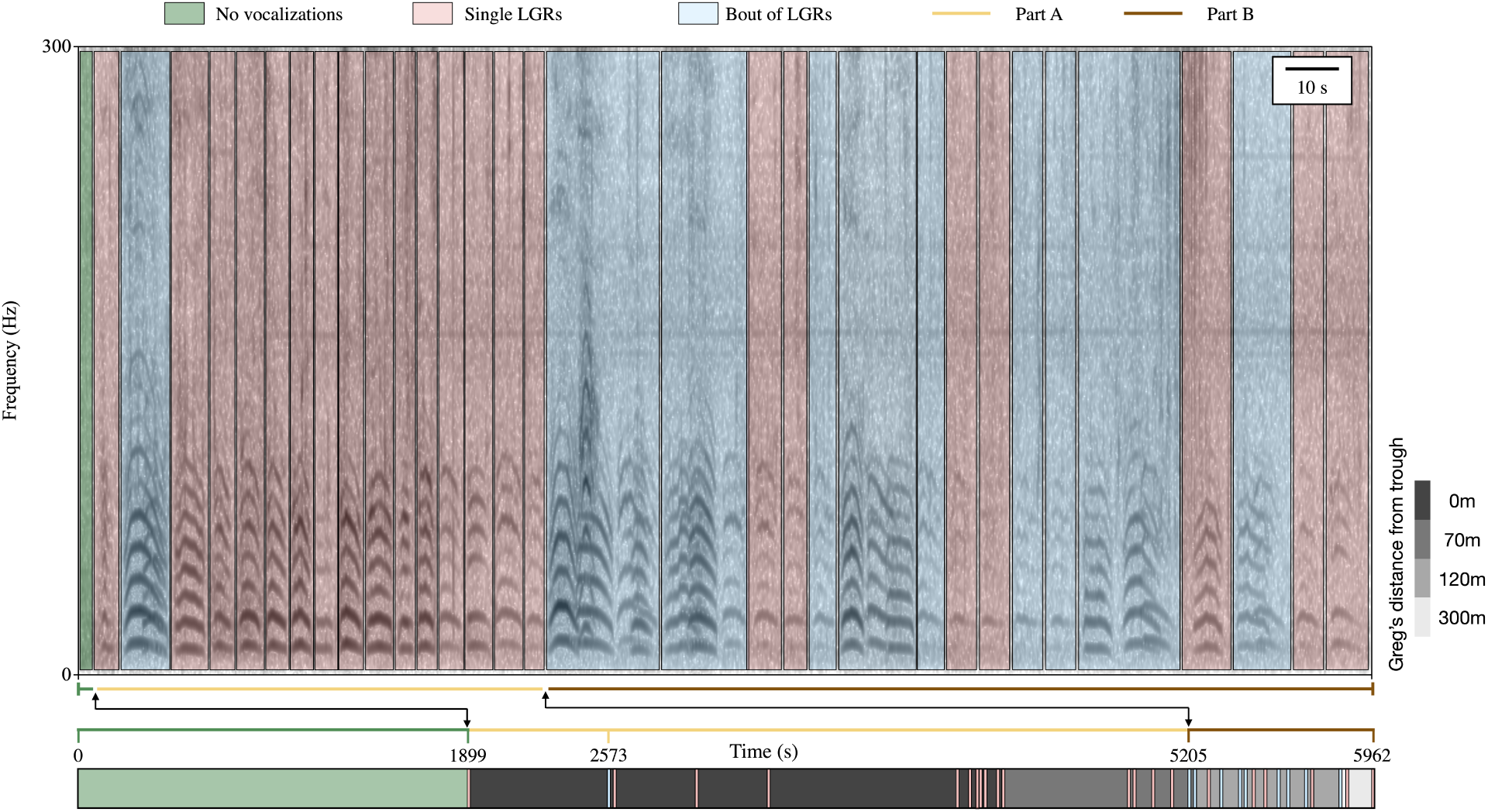
Visualization of the entirety of event 07A. Red-shaded areas indicate single, isolated rumble calls. Blue-shaded areas indicate bouts of rumbles. The time between vocalizations is removed in the main section of the plot, to show the sequence and duration of each vocalization only. The bar at the bottom contains the intervals between vocalizations to scale. Note that here the vocalizations themselves appear simply as bars of equal width. Additionally, the gray colored blocks map Greg’s distance from the trough throughout the visit, with lighter shades indicating increasing distance from the waterhole.

We have noticed that the above is a shared pattern between the waterhole visit events, that can be summarized as follows: 1) a group of bonded male elephants arrive at the waterhole, 2) there is typically an initial period with very little to no vocalization from or between the group members, 3) the dominant elephant (mostly) offers a series of single, usually unanswered rumbles and gradually starts moving away from the trough, 4) after a yet unknown trigger event, bouts of overlapping rumbles take place in a relative short period of time, 5) elephants are mobilized and leave the waterhole as a group.

We note that this behavior is of similar structure to that of the antiphonal calls among members of female elephant families [16], and we believe that it serves the same function of facilitating group coordination. Table 1 summarizes some bout event statistics. The elephant groups that were recorded and presented in this paper have up to eight members, though typically only about four of them take part in these vocalization exchanges described here. On average, each bout comprises three rumbles, and each visit event has between two to nine bouts, depending on the duration of the visit and the size of the group. It’s important to notice that the first caller in a bout is typically the leader of the group. Fig 2 shows the percentage of bouts in our dataset initiated either by the highest ranking, 2nd in rank, or any other elephant in the group.

**Figure 2:**
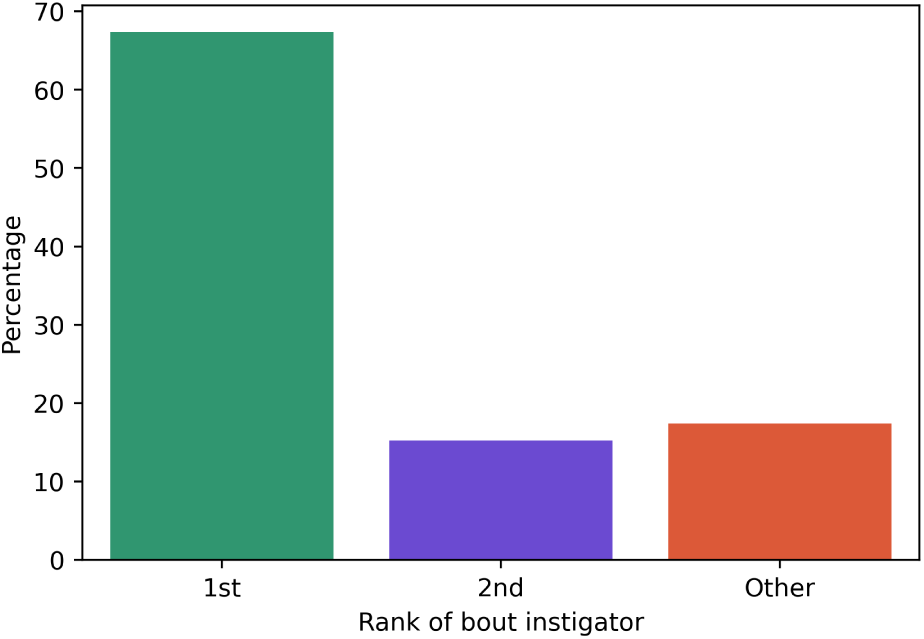
Rumble bout instigator by social rank. Percentage of bouts initiated by elephants ranked 1st, 2nd, or *other* within the social hierarchy of their groups. The majority of bouts are initiated by the no. 1 ranking elephant.

### 2.2 Frequency content of rumbles

We extracted a number of frequency features for each of the rumbles in our dataset in order to get an overall understanding of the frequency characteristics of these calls. These features are: the average fundamental frequency (*F0* ), the minimum and maximum fundamental frequency within each rumble (*minF0*, *maxF0* ), the mean slope of the F0 curve of each rumble (*meanslopeF0* ), the time during an rumble where *maxF0* occurred (*tmaxF0* ), the average of the first and the second formant (*F1*, *F2* ), and the average of the third and fourth formants (*F3* and *F4* ), in the rumbles where higher formants were present (about 80% of total number of rumbles). Here, ‘average’ is calculated over the duration of each rumble. When two rumbles overlap, we cut away the overlapping segments and the frequency features are calculated for the remaining, non-overlapping parts of the rumbles. Figs 3 and 4 show the distributions of each of these features and the pairwise relationship between them for all rumbles in our dataset. We note the bimodal distribution of the second formant F2. Most rumbles show an early increase of their fundamental frequency curve - within the first 2 seconds, though there is a smaller, yet substantial amount of rumbles where the fundamental frequency curve rises towards the end of the rumble. This finding is of course skewed by the fact that a number of rumbles have a total duration close to 2 seconds.

**Figure 3:**
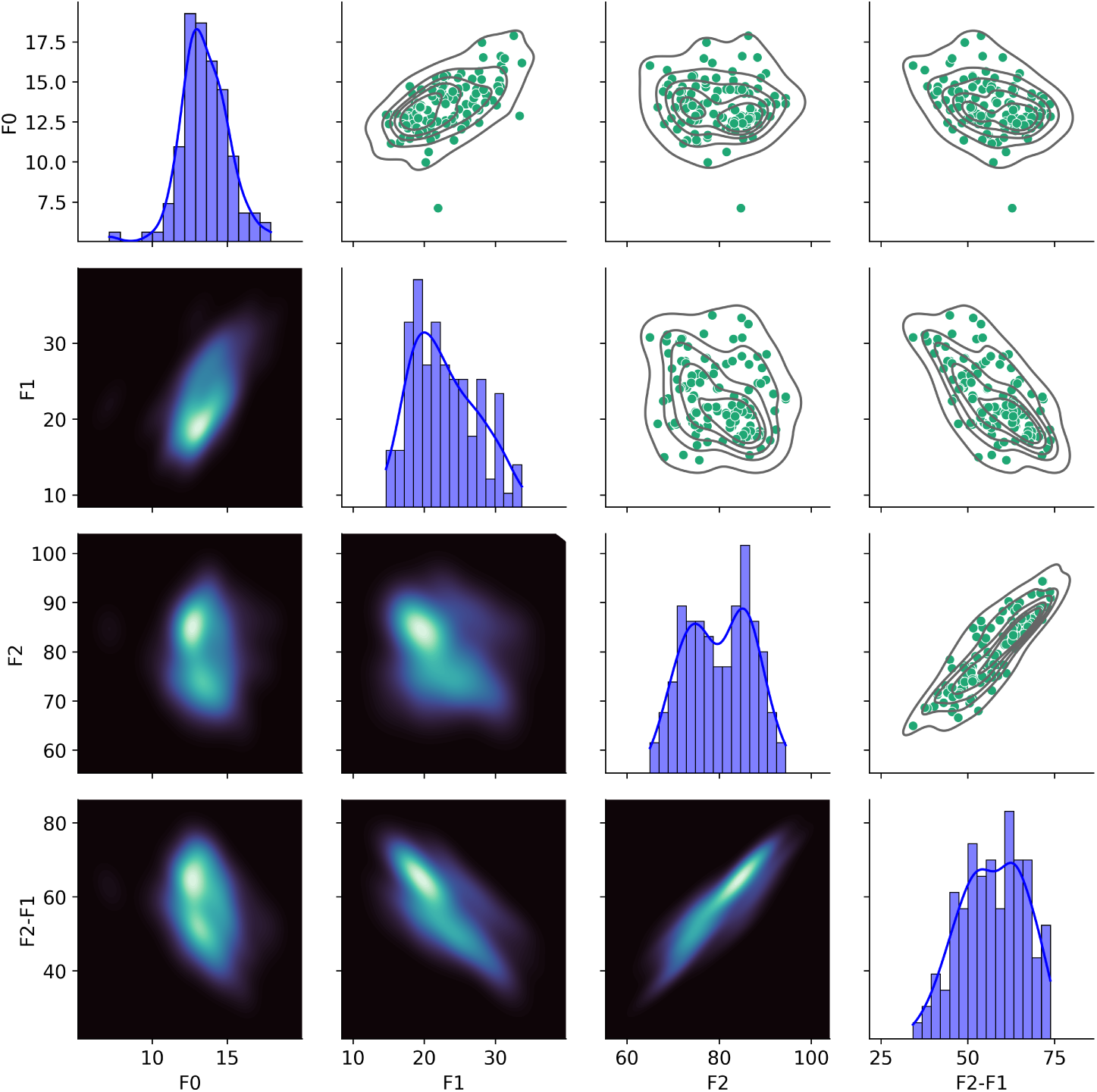
Fundamental and formant frequencies of rumbles. *Diagonal:* Distribution plots of average fundamental frequency F0, the first two formants F1 and F2, and the formant spacing F2-F1, for all the rumbles in our dataset. *Below diagonal:* Density plots of pairwise relationships between all features presented in the diagonal. *Above diagonal:* same as lower triangle but in the form of scatter plots.

**Figure 4:**
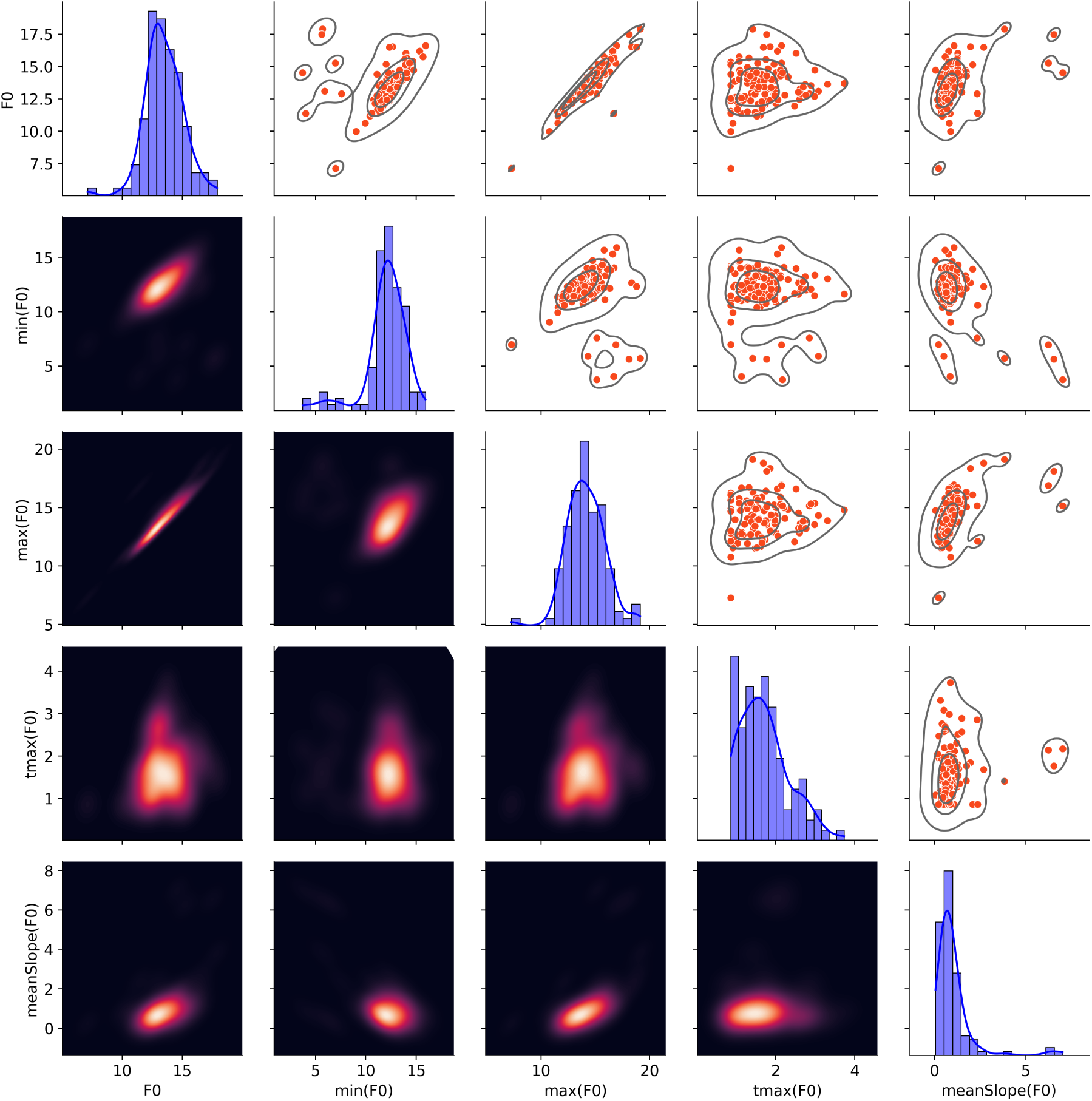
Characteristics of fundamental frequencies of rumbles. *Diagonal:* Distribution plots of average fundamental frequency F0, min and max of F0, time of maxF0, and mean slope of F0 curve for all the rumbles in our dataset. *Below diagonal:* Density plots of pairwise relationships between all features presented in the diagonal. *Above diagonal:* same as lower triangle but in the form of scatter plots.

### 2.3 Vocal spaces

In this section we take a look at how some of the rumble frequency features presented in Figs 3 and 4 vary among different individuals, within the same visit event, and across different events. We pay special attention to the first two formant frequencies, F1 and F2. The frequency plane F1-F2 is what we refer to as vocal space.

### Of individuals

Fig 5 shows the means, standard deviations, and standard error of the mean for formants F1 and F2 for each elephant. The overall formant frequency means across all rumbles in our dataset are 22.8 ± 4.6 Hz and 79.9 ± 7.1 Hz for F1 and F2 respectively, values which are in agreement with those reported in [50]. We notice that the formant means among individuals can vary significantly, although there is substantial overlap. In order to quantify whether the formant distributions we observe for each of the 9 elephants are distinct or instead originate from a common distribution, we performed Kruskal-Wallis tests for the 9 groups of F1, F2, and the formant dispersion F2-F1. The p-values returned are 3E-10, 6.8E-4, and 1.6E-8 respectively, indicating a significant difference in the formants for at least one of the 9 individuals, although it is not possible to tell from these tests alone who that elephant is, or if there are more than one whose formant frequencies distributions are statistically different from those of the rest. We address this later in this section by performing Mann-Whitney U tests.

**Figure 5:**
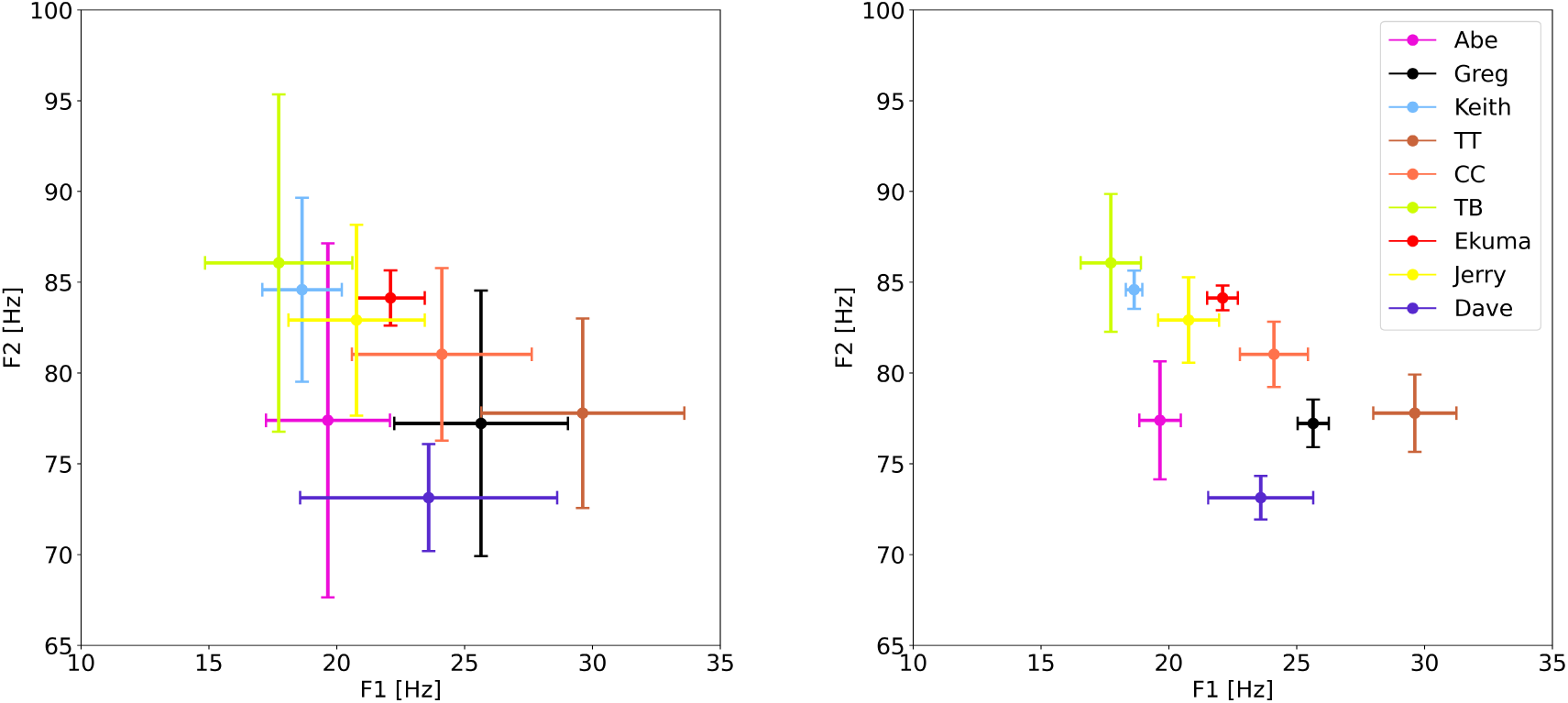
Formant mean values with SD and SEM. Mean values of the first two formants calculated from all available rumbles for each of nine elephants with *N* ≥ 5. The bars indicate Standard Deviations (SD; left) and Standard Error on the Mean (SEM; right).

The formant values used thus far are averages over the whole duration of each rumble. To take a closer look at the potential sub-structure of these calls and better represent the individual vocal space of elephants, we extracted formant values at a finer time resolution. More specifically, we divided our audio files in slices of 1000ms, 500ms, or 250ms and measured F1 and F2, for each of the slices. We noticed a definite fluctuation of the first two formants during many rumbles, but also fairly spectrally stable and flat rumbles that maintain the formant structure throughout their duration. This is a finding we will focus on in an upcoming paper. The results that follow rely on formant values calculated from sub-rumble segments of 1000ms.

Fig 6 shows the vocal spaces (i.e. the location of rumbles on the F1-F2 plane) for nine elephants. In this figure we included only elephants for which we have more than five rumbles. The figures suggest that elephants occupy overlapping yet distinct areas on the F1-F2 plane, and the Kruskal-Wallis tests mentioned earlier indicate that these areas differ statistically from each other. To locate that difference, we did pairwise Mann-Whitney U tests, to examine if the underlying formant distributions for each pair of elephants are the same or not. The tests were performed for each of the formants F1 and F2, as well as the formant dispersion F2-F1. Fig 6 shows the p-values for the test comparing the formant dispersion distributions. For 64% of the pair comparisons the p-value is below the significance level for rejecting the null hypothesis (*p <* 0.00138; see Methods for discussion on the significance level threshold). The test results for F1 and F2 are included in the Supplementary material. We notice that when comparing only the F1 or F2 distributions, in about 45-50% of the pair comparisons the null hypothesis holds.

**Figure 6:**
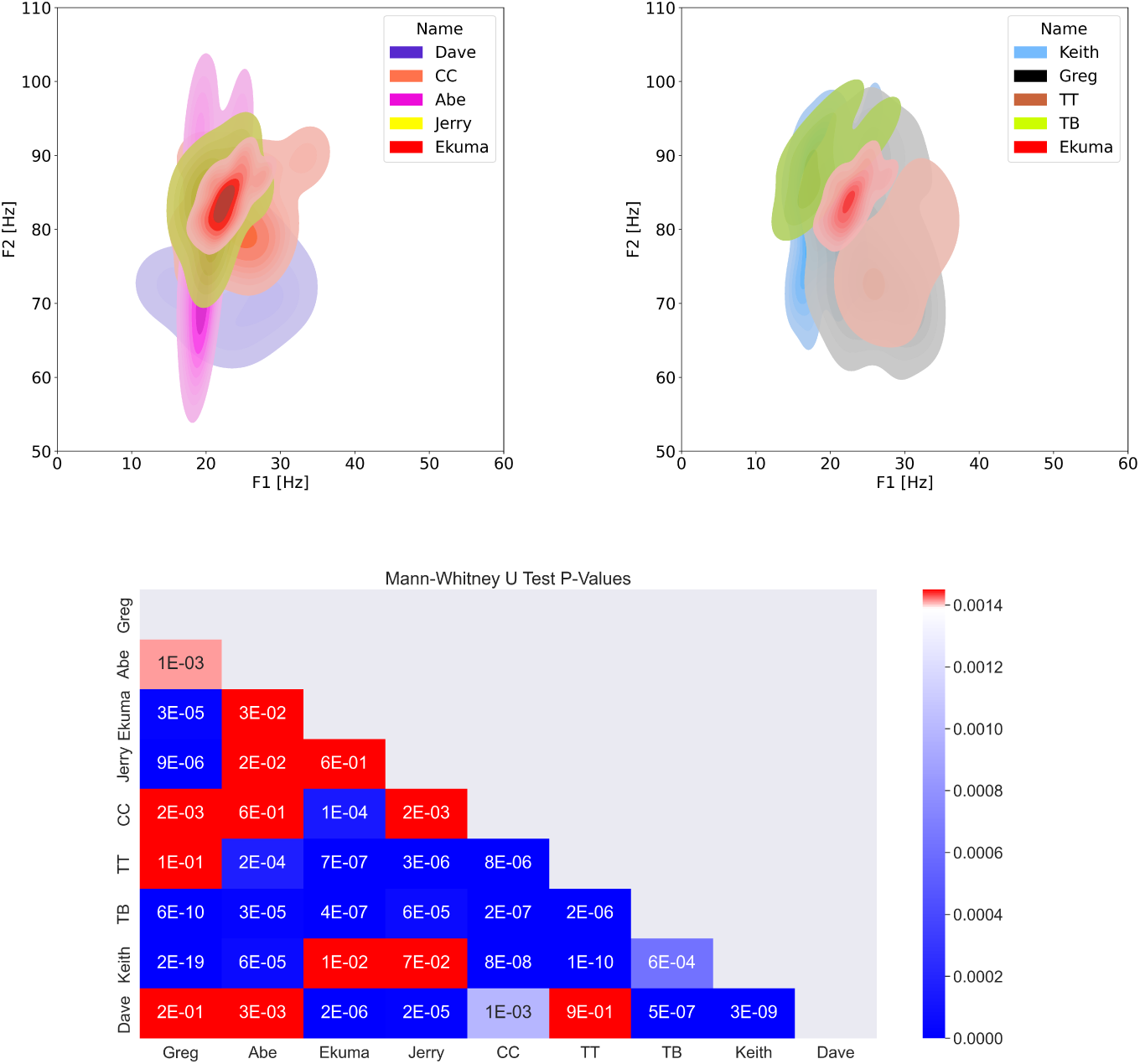
Vocal spaces of individuals elephants and pair-wise Mann Whitney U Tests. *Up:* Comparison of vocal space of nine elephants. Each panel includes five out of nine elephants, with Ekuma - in red - being present in both panels for easy visual orientation. Individuals seem to occupy overlapping yet distinct areas on the F1-F2 plane. *Down:* p-values of the Mann-Whitney U tests of the formant dispersion (*F* 2 − *F* 1) among nine individual elephants. The formant dispersion values of rumbles appear to come from different distributions for 64% of the possible pair-wise comparisons. The threshold for rejection of the null hypothesis is p-value≤0.00138 (see Methods).

In [50] the authors reported on acoustic cues in male L. Africana which hint to the identity of individuals. By using pDFA (permutated Discriminant Function Analysis) they achieved 55% accuracy when classifying by caller the rumbles of ten male elephants. We conducted a similar analysis for 9 of the elephants in our dataset. Using Principal Component Analysis (PCA) first, we kept 5 components which account for 98% of the variance in our data, and from those we identified the 5 most important of the original frequency features in our dataset. Those were: F0, F3, tmax(F0), F2, and F1. Based on those 5 features, pDFA achieved 57% accuracy in assigning the right caller to the rumbles in our dataset.

### Of events

As the members of the elephant groups in these waterhole visit events change, due to the social interaction and structure of the group over the years, we expect, given that individuals exhibit a vocal space personalization, that as a result the vocal space of different events will vary. Fig 7a shows the position of the rumbles of different visit events on the F1-F2 plane. Although overlapping, these events appear distinct.

**Figure 7:**
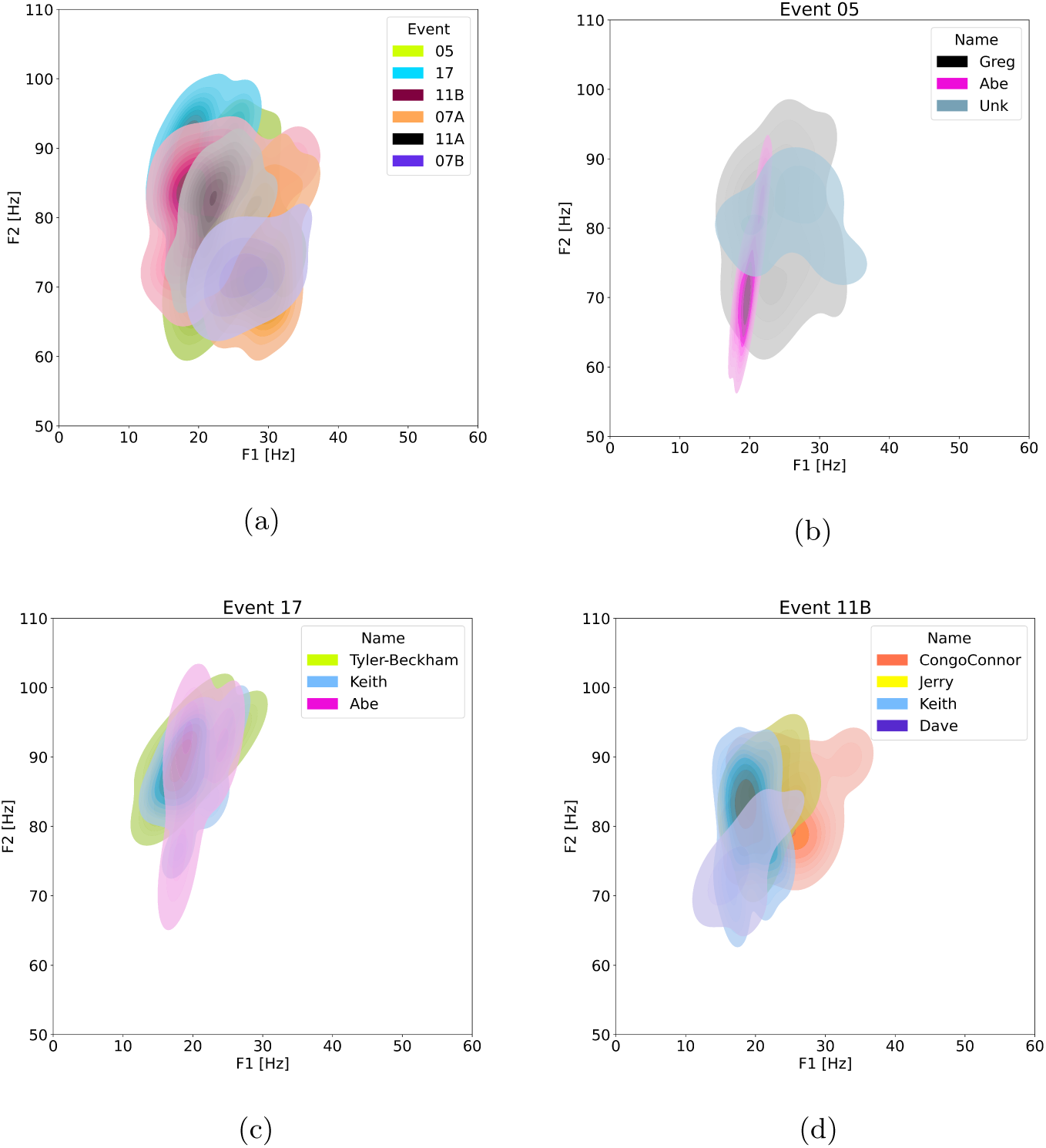
Vocal spaces of events and contribution of individuals on vocal spaces of different events. *(a)* The formant frequencies F1 and F2 of rumbles from six different waterhole visit events suggest variations in the vocal landscape of group vocalizations. *(b)-(d)*Vocal spaces of three of the events shown in panel *(a)* are deconstructed into vocal spaces of the individual callers in those events.

### Of individuals in events

The plot and analysis results showing individuality of rumbles considered all rumbles for each individual, from all events during the years. To infer whether individuals alter their vocal space between events, we examine how the vocal space of individuals vary from one event to the other. We also compare the vocal spaces of different individuals who partake in the same event. Fig 7 shows the event-specific vocal spaces of elephants who appear in more than one waterhole visits in our dataset. Fig 8 compares the vocal spaces of different group members within the same event, for two distinct events.

**Figure 8:**
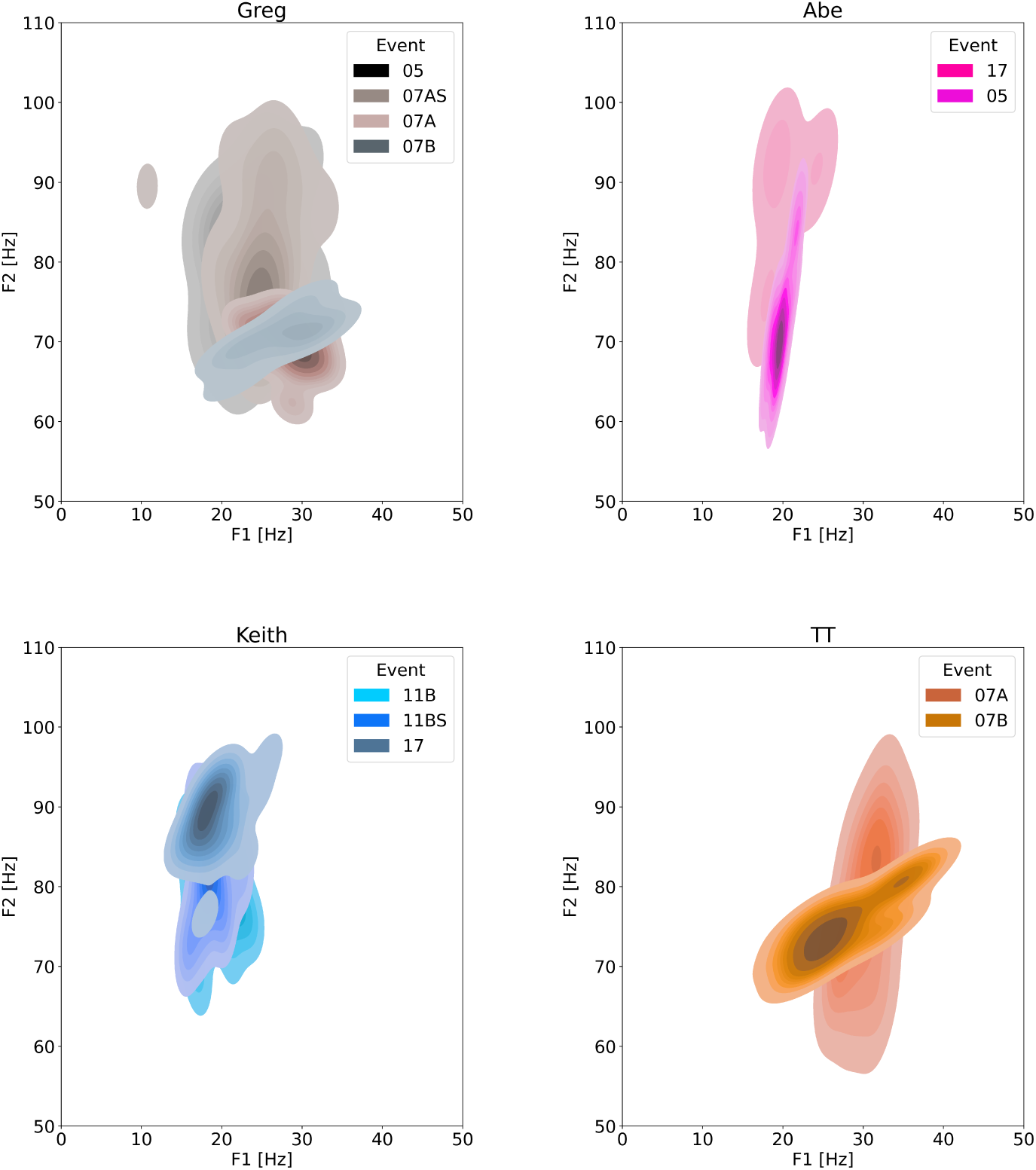
Comparison of personal vocal spaces across different events. The vocal spaces of elephants Greg, Abe, Keith, and TT during various events. Individuals appear to sometimes vary their vocal space from event to event. Social context, aging, and information content of the vocal exchanges might be contributing factors to this variation.

### The role of aging

To examine whether the change of an individual’s vocal space might be attributed to aging, we compared rumbles of individuals for whom we had data far apart in time to showcase a potential aging effect. Fig 9 shows the formant distance (F2-F1) and fundamental frequency (F0) of rumbles for 4 elephants. For three of the elephants there are data from two different events with a time gap between four to 12 years. The data for Greg are only two years apart and are presented as a counterexample. Abe, Dave, and Keith show an increase of F2-F1 from the earlier to the later event. More specifically Abe, a full adult during the whole study period, shows an increase in formant distance from 2005 to 2017, although the fundamental frequencies remain the same, indicative of no change in the size of Abe’s vocal chords. Examining separately F2 and F1 for the 2005 and 2017 events, we notice that the increase in the formant spacing is due to the increase of F2 (F1 remains stable). Keith, a young adult in 2011, but fully matured in 2017, exhibits an obvious increase of F2 with F1 remaining unaltered. His fundamental frequency also does not show any noticeable change. Dave transitioned from adolescence to early adulthood around the time of his earlier rumbles in 2007. Although we have less data for him, his later rumbles from 2011 show higher formant distance, but in his case this is a result of lower F1 frequencies and only somewhat elevated F2 frequencies. His fundamental frequencies also remain unchanged. Greg is also included in this plot, but we would not expect any aging effects in his case as the data are only 2 years apart and he was already a fully grown elephant. We notice an inverse trend in his case: the formant distances in his rumbles decrease in the 2007 event compared to the 2005 event. This is a result of a simultaneous increase of F1 and decrease of F2, quite unlikely to what we observed for the three other elephants. His fundamental frequencies also remain stable throughout.

**Figure 9:**
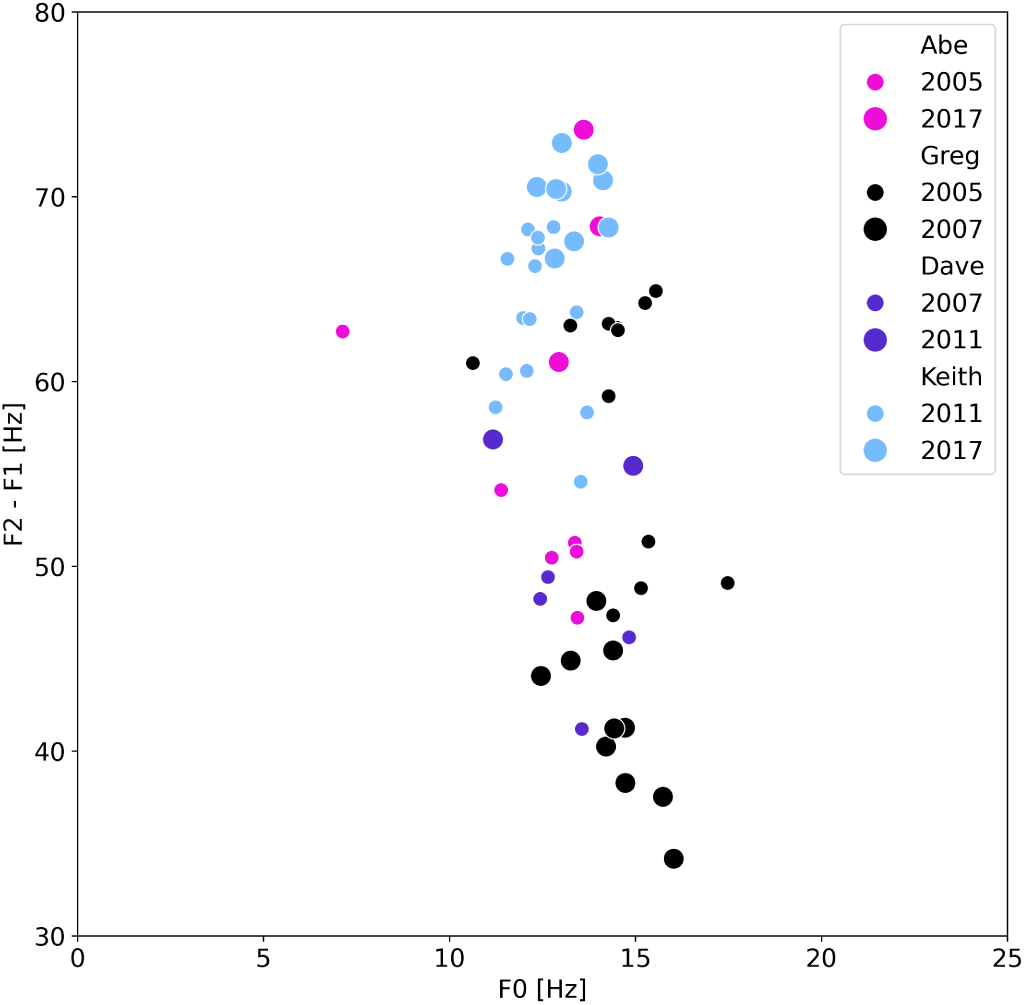
Development of fundamental and formant frequencies over time. Formant distance and fundamental frequencies for 4 elephants. For each elephant data are from two different years denoted by different symbol sizes.

## 3 Discussion

In this paper, we documented for the first time the use of antiphonal calls in wild male African elephants in the context of group departure coordination. The dynamics we described mirror the antiphonal calls observed in female elephant families, suggesting an equivalent role of turn-taking and rumbles in coordinating group actions and facilitating group decisions. We also presented novel findings which highlight that elephant vocal spaces reflect personalized traits and can be influenced by social context.

The turn-taking rumble exchanges we presented include partial overlaps between consecutive calls. 59% of rumbles that were followed by another rumble (i.e. rumbles that were not the last call in a bout), were overlapped by the succeeding rumble. The average overlapping time was 1.5s, whereas the average duration of a rumble was 5s. It has been suggested that in birds, overlapping might in itself be a signal [51].

In a study of turn-taking vocal exchanges in bonobos and how those exchanges drive social bonding [28], the authors remark on some findings which share remarkable commonalities with some of ours. Like the elephants in our study, bonobos exhibit three call patterns: i) single, isolated calls, ii) successive calls (*<* 2500ms inter-call durations) with no overlap, and iii) overlapping calls. Group turn-taking calls in bonobos usually include two to three callers. The average number of callers in the rumble bouts we presented is three to five depending on the group size. Bonobos utter 3.7 (±2.1) calls per interaction. The elephants in our dataset average 3.1 rumbles per bout. Finally, socially affiliated bonobos are preferred vocal partners, a finding that holds true for the elephants in our study and those of [40] and [41]. Further insights in the role of affiliation in vocalization exchanges of male L. Africana will be presented in an upcoming paper.

We have noticed the following features in the turn-taking rumble exchanges:

1. The order of the speaker turns varies, although 60% of the bouts are initiated by the highest ranking elephant. 2) The number of participants in an event can vary. 3) Predominantly, only one elephant vocalizes at a time. 4) Overlapping does exist, as reported in the Results section, but it takes place sometimes and with - what seems to be - designed overlapping period. 5) The turn size is not fixed, but rather rumbles have quite a wide range of duration (the average duration of a rumble is about 5s, with values ranging from just below 2s all the way up to 8s or 9s). The above match five out of eight of Sacks’ rules [44] that govern turn-taking in human conversations. The remaining three rules refer to qualities and quantities related to spontaneity of the speaker’s content and ways to regulate and repair turn exchange errors, something that is not accessible for these rumble calls, either due to lack of the right data or impossibility - at least for now - to access the information content of these vocal exchanges.

In our investigation of vocal spaces, we have shown that individuals exhibit statistically unique positions within the F1-F2 frequency plane. Moreover, the vocal space of an elephant may differ across various events, and the collective vocal space of a group of elephants can also vary depending on the group’s membership composition. The difference in the vocal space of individuals that we observed and analyzed here, might be attributed to a variety of factors. Those could be biological/anatomical factors like age or size of individuals. It has been shown that formant frequencies of ring-tailed lemurs carry information of individual identity, predominantly due to morphological variations of the shape rather than the length of the vocal tract [52]. It is very likely that the individuality of the elephant vocal spaces we have presented here, is a result of similar anatomical variability in elephant vocal tracts. Age however, did not seem to have a consistent effect on the vocal spaces of 4 individuals, at least not in the anticipated way based on the assumed vocal fold and vocal tract length increase with increasing age.

Vocal space differentiation could also reflect a personal signature, which could depend on the particulars of the event, like the composition of the group, the role or rank of individuals within it, etc. In an interesting study on social rumbles of wild mature male L. Africana [49], the authors showcased that elephants produced unique vocalizations facilitating individual recognition. They identified 5 spectral features that contributed almost evenly to these personalized variations. Although they mentioned that the use of formant frequencies was technically not possible in their analysis, two of the spectral features used (center frequency and frequency 95%) have mean values close to the ranges of F1 and F2 values that we have measured in our study of rumbles.

We furthermore believe that the vocal space differentiation we presented in this study encodes semantic information, akin to the formant frequency variation in vowels of human languages. However, further analysis on the type and amount of the information content of these vocal exchanges is required to establish such a finding. We will expand our inquiry towards that direction in an upcoming paper.

We posit that elephant vocal communication, particularly in regards to turn-taking dynamics and the characteristics and role of overlapping vocalizations and associated vocal spaces, warrants further targeted investigation. This exploration could be pivotal for unraveling the intricacies of elephant vocal generation and vocal structural composition, as well as language processing mechanisms. Additionally, in-depth analysis of these communicative calls can yield profound insights into the contextual and informational aspects of elephant vocal interactions. This is a rich area for future research, where significant discoveries on elephant communication and social dynamics and behavior await.

## 4 Methods

The analyses reported here were conducted not on the raw recordings themselves, but on a reduced dataset of derived acoustic and spectral features (see Section 4).

### Raw Data

The raw audio data were recorded during the period 2005-2017 at Dr. Caitlin O’Connell’s long-term field site at a remote permanent waterhole at Namibia’s Etosha National Park. Recordings came with labels including the date/time and the name of the vocalizer, and, for some of them, the location of the individuals relative to the waterhole and the microphone. Details on the observation site, recording conditions, identification of individuals etc., can be found in previous publications [16, 53].

### Data Description

The reduced dataset used for analysis contains rumbles from six different waterhole visit events. For two of these events (07AS and 11BS), the dataset also includes a ”prelude” sequence of single, unanswered rumbles produced by the groups highest-ranking elephant. The number of rumbles, participating individuals, and event details are provided in Table 1.

### Data Reduction and Analysis

The reduced dataset was generated from the raw audio recordings by extracting acoustic and spectral features using Praat [54], a free software package for speech analysis. Praat’s scripting functionality was employed to automate and expedite the extraction process. Frequency features - including formant values and fundamental frequency - were computed from the original WAV files. Praat was also used for visual inspection of spectrograms and spectral plots.

All subsequent data handling, statistical analyses, and figure generation were performed in Python.

#### Extraction of spectral features with Praat

Attempting to use Praat with its default settings to extract formant values or formant trajectories for elephant rumbles is understood to give unreliable results. To obtain accurate results, specific parameters must be set beyond the softwares defaults and verified meticulously using a substantial number of vocalization data. This necessity applies broadly to various types of non-human animal vocalisations. The procedure we employed to ensure reliable formant and fundamental frequency extraction using Praat is detailed below. The scripting (via Praat) of the process described below was done for the purpose of reproducibility and automating this procedure for handling large number of files.

Importantly, this process allows us to use the same parameter settings for all file treatment, therefore minimising human bias.

#### For formant frequencies extraction

1. Each audio file is resampled to 1000Hz, giving a Nyquist frequency of 500Hz. Reasoning: visual inspection of all 120 audio files have shown that there is no power above 500Hz (in reality, from a little below 500Hz), even for rumbles that clearly show 3rd and 4th formants. Note: the Precision parameter used for the interpolation during resampling is set to 50 samples.
2. The spectrum of the resampled audio file is calculated (zero padding for Fast Fourier Transform selected)
3. LPC (linear predictive coding) smoothing on the spectrum from the previous step, using 9 peaks and pre-emphasis from 50 Hz.
4. Formant peaks are then identified by Praat from the above smoothed spectrum, with the maximum number of possible peaks set to 5.

#### For fundamental frequencies extraction

1. A band-pass filter (Hann window) is applied from 0-1000Hz and smoothing of 10Hz.
2. The fundamental (pitch) trajectory is calculated using To Pitch (ac), with customised settings as follows: Time step = 0.01s, Pitch floor=3.5Hz, Max number of candidates = 2, Very accurate = ”yes”, silent threshold = 0.03, voicing threshold = 0.45, octave cost = 0.1, octave jump cost = 0.35, voiced/unvoiced cost = 0.14, Pitch ceil = 35Hz
3. From the trajectory calculated above, we extract (with Praat) the following quantities: mean value, standard deviation, maximum value, minimum value, mean slope of the trajectory, time, i.e., location of maximum F0 value. Parabolic interpolation is used in calculating the above values.

The parameters chosen for this process (the number of peaks for LPC, pre-emphasis limit, time steps, pitch limits, etc) were optimised through careful, manual trials to ensure the resulting formant (and fundamental) frequency values match those extracted via visual examination of the spectra from approximately 20 audio files. Once the reliability of these parameters was established, we scripted the procedure in Praat and applied it to all audio files (about 120 in total). Scripting allows control over a larger number of parameters than does the GUI, which is adapted to human vocalisations. The resulting formants for each file were then validated through visual inspection of each spectrum. This same procedure, on the same files, was repeated a couple more times: Praat script runs on all audio files, automatically extracting formants for each one (average over full duration), and then those values are confirmed manually (visual inspection) by identifying the formant bumps (and fundamental sharp peaks) in the spectrum for each of the 120 files.

#### Vocal spaces

We use the non-parametric Kruskal-Wallis test instead of ANOVA, as our data do not fulfill the requirement for homoscedasticity. Heteroscedasticity was confirmed by running a Levene test.

The extraction of the fundamental frequency’s curve and higher formants (3rd formant or higher) is harder when we use smaller audio segments, and requires manual inspection of the resulting values to confirm or correct. This gets more challenging the smaller the audio segments become as the number of files to extract formants from increases. We decided to only extract F1 and F2 for our sub-rumble investigation frequency features extraction, as those two frequencies are more robust and we can use a script to automatically perform the task. For the Mann-Whitney U tests, as multiple hypotheses are tested simultaneously, we adjust the significance level for rejecting the null hypothesis (typically p-value*<* 0.05) according to the Bonferonni correction. The corrected p-value threshold is thus set to 0.05*/m* = 0.0014, where *m* = 36 is the number of tested hypotheses, i.e. unique pairs of elephants in our dataset.

## Acknowledgements

The raw data of elephant rumbles were provided by Dr. Caitlin O’Connell. E.R. would like to thank Prof. L. Mahadevan for hosting her in the SoftMath Lab during the preparation of the work included in this manuscript. E.R. would like to acknowledge funding from the Department of Systems Biology at Harvard Medical School.

## Supplementary material

### Reduced data

The reduced data used for this study - comprising acoustic features such as formant and fundamental frequencies - can be accessed at the following repository: https://github.com/emmarant/elephant_vocalizations/tree/master/rumbles

### Extra plots

**Figure 10:**
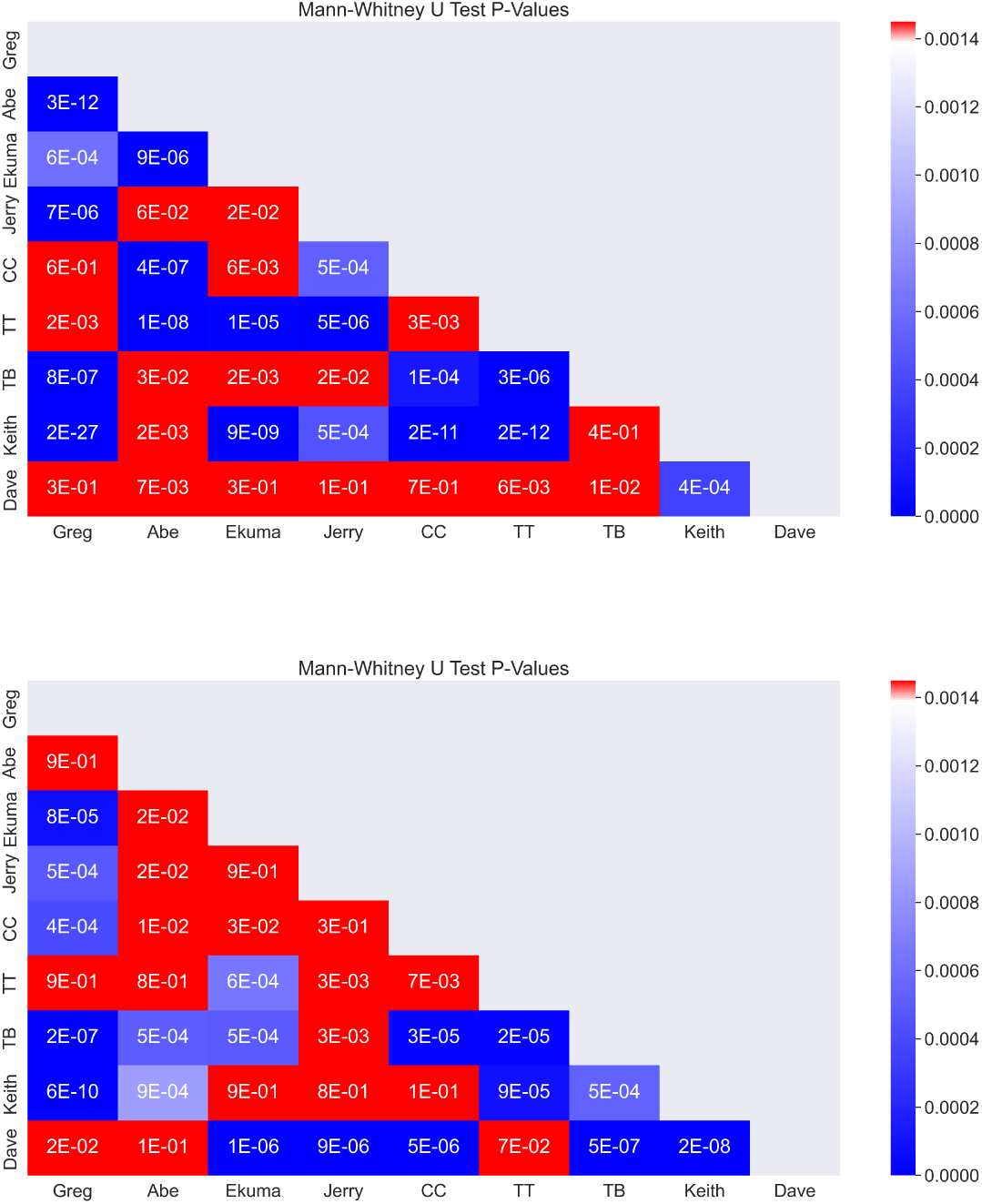
Pair-wise Mann-Whitney U tests for F1 and F2. p-values of the Mann-Whitney U tests of the formant frequencies F1 (up) and F2 (down) among nine individual elephants. The values of the first two formants appear to come from different distributions for 50% of the possible pair-wise comparisons. The threshold for rejection of the null hypothesis is p-value≤0.00138 (see Methods).

## References

[1] Jack W. Bradbury and Sandra L. Vehrencamp. *Principles of animal communication*. Sinauer Associates, Sunderland, MA, 1998.

[2] Claudia Fichtel and Marta Manser. *Vocal communication in social groups*, pages 29–54. Springer Berlin Heidelberg, 2010.

[3] Rianna Burnham. Animal calling behaviours and what this can tell us about the effects of changing soundscapes. Acoustics, 5:631–652, 7 2023.

[4] Rogério Grassetto Teixeira da Cunha and Richard W. Byrne. The Use of Vocal Communication in Keeping the Spatial Cohesion of Groups: Intentionality and Specific Functions, pages 341–363. Springer New York, 2009.

[5] Larissa Conradt and Timothy J. Roper. Group decision-making in animals. Nature, 421:155–158, 1 2003.

[6] Larissa Conradt and Timothy J. Roper. Consensus decision making in animals. Trends in Ecology & Evolution, 20:449–456, 8 2005.

[7] Sue Boinski. The coordination of spatial position: a field study of the vocal behaviour of adult female squirrel monkeys. Animal Behaviour, 41:89–102, 1 1991.

[8] Hans Kummer. Social organization of hamadryas baboons. A field study. Bibliotheca primatologica no. 6. Karger, Basel ; New York, 1968.

[9] Sue Boinski. Vocal coordination of troop movement among whitefaced capuchin monkeys, cebus capucinus. American Journal of Primatology, 30:85–100, 1 1993.

[10] Sue Boinski and Aimee F. Campbell. Use of trill vocalizations to coordinate troop movement among white-faced capuchins: a second field test. Behaviour, 132:875–901, 1995.

[11] Anna Lucia Sperber, Lynne M. Werner, Peter M. Kappeler, and Claudia Fichtel. Grunt to govocal coordination of group movements in redfronted lemurs. Ethology, 123:894–905, 12 2017.

[12] Alexander H. Harcourt and Kelly J. Stewart. Gorillas’ vocalizations during rest periods: Signals of impending departure? Behaviour, 130(1-2):29–40, 1994.

[13] Jeffrey M. Black. Preflight signalling in swans: A mechanism for group cohesion and flock formation. Ethology, 79:143–157, 1 1988.

[14] Reena H. Walker, Andrew J. King, J. Weldon McNutt, and Neil R. Jordan. Sneeze to leave: African wild dogs ( lycaon pictus ) use variable quorum thresholds facilitated by sneezes in collective decisions. Proceedings of the Royal Society B: Biological Sciences, 284:20170347, 9 2017.

[15] Joyce H. Poole, Katherine Payne, William R. Langbauer, and Cynthia J. Moss. The social contexts of some very low frequency calls of african elephants. Behavioral Ecology and Sociobiology, 22:385–392, 6 1988.

[16] C. E. O’Connell-Rodwell, J. D. Wood, M. Wyman, S. Redfield, S. Puria, and L. A. Hart. Antiphonal vocal bouts associated with departures in free-ranging african elephant family groups ( loxodonta africana ). Bioacoustics, 21:215–224, 10 2012.

[17] T. K. Athira and T. N. C. Vidya. Elephant social systems: What do we know and how have molecular tools helped? Journal of the Indian Institute of Science, 101:257–278, 4 2021.

[18] Iain Douglas-Hamilton. On the ecology and behaviour of the african elephant. *PhD Thesis,* University of Oxford, 1972.

[19] Cynthia Moss and Joyce Poole. Relationships and social structure in African elephants, pages 315–325. 01 1983.

[20] R. M. Laws and I. S. C. Parker. Recent studies on elephant populations in east africa. Symposia of the Zoological Society of London, 21:319–359, 1968.

[21] George Wittemyer, Iain Douglas-Hamilton, and Wayne M. Getz. The socioecology of elephants: analysis of the processes creating multitiered social structures. Animal Behaviour, 69:1357–1371, 6 2005.

[22] Joyce H. Poole. Rutting behavior in african elephants: the phenomenon of musth. Behaviour, 102:283–316, 1987.

[23] Judith Kay Berg. Vocalizations and associated behaviors of the african elephant ( loxodonta africana ) in captivity. Zeitschrift fr Tierpsychologie, 63:63–79, 1 1983.

[24] K. M. Leong, A. Ortolani, K. D. Burks, J. D. Mellen, and A. Savage. Quantifying acoustic and temporal characteristics of vocalizations for a group of captive african elephant loxodonta africana. Bioacoustics, 13:213– 231, 1 2003.

[25] Joseph Soltis. Vocal communication in african elephants ( loxodonta africana ). Zoo Biology, 29:192–209, 3 2010.

[26] Patrick J. Clemins, Michael T. Johnson, Kirsten M. Leong, and Anne Savage. Automatic classification and speaker identification of african elephant ( loxodonta africana ) vocalizations. The Journal of the Acoustical Society of America, 117:956–963, 2 2005.

[27] Andrea Ravignani, Laura Verga, and Michael D. Greenfield. Interactive rhythms across species: the evolutionary biology of animal chorusing and turntaking. Annals of the New York Academy of Sciences, 1453:12–21, 10 2019.

[28] Florence Levréro, Sonia Touitou, Julia Frédet, Baptiste Nairaud, Jean-Pascal Gury, and Alban Lemasson. Social bonding drives vocal exchanges in bonobos. Scientific Reports, 9:711, 1 2019.

[29] Marlen Fröhlich, Paul Kuchenbuch, Gudrun Müller, Barbara Fruth, Takeshi Furuichi, Roman M. Wittig, and Simone Pika. Unpeeling the layers of language: Bonobos and chimpanzees engage in cooperative turn-taking sequences. Scientific Reports, 6:25887, 5 2016.

[30] Noriko Katsu, Kazunori Yamada, Kazuo Okanoya, and Masayuki Nakamichi. Temporal adjustment of short calls according to a partner during vocal turn-taking in japanese macaques. Current Zoology, 65:99–105, 2 2019.

[31] Cecilia P. Chow, Jude F. Mitchell, and Cory T. Miller. Vocal turn-taking in a non-human primate is learned during ontogeny. Proceedings of the Royal Society B: Biological Sciences, 282:20150069, 5 2015.

[32] Daniel Y. Takahashi, Darshana Z. Narayanan, and Asif A. Ghazanfar. Coupled oscillator dynamics of vocal turn-taking in monkeys. Current Biology, 23:2162–2168, 11 2013.

[33] A. Lemasson, L. Glas, S. Barbu, A. Lacroix, M. Guilloux, K. Remeuf, and H. Koda. Youngsters do not pay attention to conversational rules: is this so for nonhuman primates? Scientific Reports, 1:22, 6 2011.

[34] David Symmes and Maxeen Biben. *Conversational Vocal Exchanges in Squirrel Monkeys*, pages 123–132. Springer Berlin Heidelberg, 1988.

[35] Tadamichi Morisaka, Yayoi Yoshida, Yuichiro Akune, Hideki Mishima, and Sayo Nishimoto. Exchange of signature calls in captive belugas (delphinapterus leucas). Journal of Ethology, 31:141–149, 5 2013.

[36] Tyler M. Schulz, Hal Whitehead, Shane Gero, and Luke Rendell. Overlapping and matching of codas in vocal interactions between sperm whales: insights into communication function. Animal Behaviour, 76:1977–1988, 12 2008.

[37] P. J. O. Miller, A. D. Shapiro, P. L. Tyack, and A. R. Solow. Call-type matching in vocal exchanges of free-ranging resident killer whales, orcinus orca. Animal Behaviour, 67:1099–1107, 6 2004.

[38] Nicola J. Quick and Vincent M. Janik. Bottlenose dolphins exchange signature whistles when meeting at sea. Proceedings of the Royal Society B: Biological Sciences, 279:2539–2545, 7 2012.

[39] Vlad Demartsev, Ariana Strandburg-Peshkin, Michaela Ruffner, and Marta Manser. Vocal turn-taking in meerkat group calling sessions. Current Biology, 28:3661–3666.e3, 11 2018.

[40] Joseph Soltis, Kirsten Leong, and Anne Savage. African elephant vocal communication i : antiphonal calling behaviour among affiliated females. Animal Behaviour, 70:579–587, 9 2005.

[41] Katherine A. Leighty, Joseph Soltis, Kirsten Leong, and Anne Savage. Antiphonal exchanges in african elephants (loxodonta africana): Collective response to a shared stimulus, social facilitation, or true communicative event? Behaviour, 145:297–312, 2008.

[42] Caitlin O’Connell. Elephant Don : the politics of a pachyderm posse. The University of Chicago Press, Chicago, 2015.

[43] Simone Pika, Ray Wilkinson, Kobin H. Kendrick, and Sonja C. Vernes. Taking turns: bridging the gap between human and animal communication. Proceedings of the Royal Society B: Biological Sciences, 285:20180598, 6 2018.

[44] Harvey Sacks, Emanuel A. Schegloff, and Gail Jefferson. A simplest systematics for the organization of turn-taking for conversation. Language, 50:696–735, 1974.

[45] Mary Catherine Bateson. Mother-infant exchanges: The epigenesis of conversational interaction. Annals of the New York Academy of Sciences, 263:101–113, 9 1975.

[46] Stephen C. Levinson. Turn-taking in human communication origins and implications for language processing. Trends in Cognitive Sciences, 20:6–14, 1 2016.

[47] Katharine B. Payne, William R. Langbauer, and Elizabeth M. Thomas. Infrasonic calls of the asian elephant (elephas maximus). Behavioral Ecology and Sociobiology, 18:297–301, 2 1986.

[48] Amy MorrisDrake and Hannah S. Mumby. Social associations and vocal communication in wild and captive male savannah elephants loxodonta africana. Mammal Review, 48:24–36, 1 2018.

[49] Kaja Wierucka, Michelle D. Henley, and Hannah S. Mumby. Acoustic cues to individuality in wild male adult african savannah elephants ( loxodonta africana ). PeerJ, 9:e10736, 1 2021.

[50] Angela S. Stoeger and Anton Baotic. Information content and acoustic structure of male african elephant social rumbles. Scientific Reports, 6:27585, 6 2016.

[51] Marc Naguib and Daniel J. Mennill. The signal value of birdsong: empirical evidence suggests song overlapping is a signal. Animal Behaviour, 80:e11– e15, 9 2010.

[52] Marco Gamba, Livio Favaro, Alessandro Araldi, Valentina Matteucci, Cristina Giacoma, and Olivier Friard. Modeling individual vocal differences in group-living lemurs using vocal tract morphology. Current Zoology, 63:467–475, 8 2017.

[53] C.E. O’Connell-Rodwell, J.D. Wood, C. Kinzley, T.C. Rodwell, C. Alarcon, S.K. Wasser, and R. Sapolsky. Male african elephants ( loxodonta africana ) queue when the stakes are high. Ethology, Ecology, and Evolution, 23:388–397, 10 2011.

[54] Paul Boersma and David Weenink. Praat: doing phonetics by computer. http://www.praat.org.

